# Profiling *Myxococcus xanthus* swarming phenotypes through mutation and environmental variation

**DOI:** 10.1101/2021.06.22.449537

**Authors:** Linnea J. Ritchie, Erin M. Curtis, Kimberly A. Murphy, Roy D. Welch

**Affiliations:** Department of Biology, Syracuse University, Syracuse, NY, USA; Department of Biology, Augustana College, Rock Island, Il, USA

**Author notes:** Address correspondence to Roy D. Welch.

## Abstract

*Myxococcus xanthus* is a bacterium that lives on surfaces as a predatory biofilm called a swarm.As a growing swarm feeds on prey and expands, it displays dynamic multicellular patterns such as traveling waves called ripples and branching protrusions called flares. The rate at which a swarm expands across a surface, and the emergence of the coexisting patterns, are all controlled through coordinated cell movement. *M. xanthus* cells move using two motility systems known as Adventurous (A) and Social (S). Both are involved in swarm expansion and pattern formation. In this study, we describe a set of *M. xanthus* swarming genotype-to-phenotype associations that include both genetic and environmental perturbations. We identified new features of the swarming phenotype; recorded and measured swarm expansion using time-lapse microscopy; and compared the impact of mutation on different surfaces. These observations and analyses have increased our ability to discriminate between swarming phenotypes and provided context that allowed us to identify some phenotypes as improbable ‘outliers’ within the *M. xanthus* swarming phenome.

**Importance:** *Myxococcus xanthus* grows on surfaces as a predatory biofilm called a swarm. A feeding swarm expands by moving over and consuming prey bacteria. In the laboratory, a swarm is created by spotting cell suspension onto nutrient agar in lieu of prey. The cells quickly settle on the surface and the new swarm then expands radially. An assay that measures the expansion rate of a swarm of mutant cells is the first, and sometimes only, measurement used to decide whether a particular mutation impacts swarm motility. We have broadened the scope of this assay by increasing the accuracy of measurements and reintroducing prey, resulting in new identifiable and quantifiable features that can be used to improve genotype-to-phenotype associations.

## Introduction

*Myxococcus xanthus* is a Gram-negative bacterium that moves across surfaces as a motile biofilm called a swarm. In the laboratory, swarm movement is almost always studied on an agar surface. When several million *M. xanthus* cells in a few microliters of liquid suspension are spotted onto a nutrient agar plate, the liquid is absorbed into the agar within a few minutes, leaving the cells settled on the surface in the shape of a circle. After some time, the cells begin to move, and the circle becomes a swarm. Under permissive conditions, a swarm will expand outward in all directions and double or triple in diameter within a few days. (1–3)

Swarming *M. xanthus* cells move using two mechanistically distinct and genetically separable molecular motors. These drive two different types of surface movement called Adventurous (A) motility and Social (S) motility respectively (4, 5). Cells lacking both motors do not move on agar, and swarms of such cells do not expand. The A motor functions within a cell by forming points of adhesion to an external surface, such as agar, then rolling the cell like a tank tread or twisting it like a threaded screw (6–8). The S motor functions by extending a fiber called a pilus from one end of the cell, tethering the pilus to an external object, and then retracting it like a winch (9–12). The A and S motors differ in that the A motor seems to involve a pushing force, whereas the S motor involves a pulling force. Mutant swarms possessing only the A motor (A+S-) expand faster on harder agar than swarms possessing only the S motor (A-S+), whereas the opposite is true on softer agar (4, 5, 13). Perhaps hard agar allows the A motility-associated adhesion points to more easily gain a purchase against which the cell must roll or twist itself forward, while soft agar allows the S motility-associated pili to more easily winch the cell forward because of its lower drag coefficient. Both A and S motors periodically reverse direction, swapping the leading and lagging poles of each rod-shaped cell.

This mechanistic description of A and S motors provides a framework for the interpretation of new data regarding the impact of mutation on swarm motility. Specifically, the expansion rate on agar of a swarm of mutant cells (which we refer to as a mutant swarm) is compared to wild type (WT) to measure the mutation’s phenotypic impact with respect to motility. Further, a comparison of that swarm’s relative expansion rates on hard and soft agar provides an indication of how much the A and S motors, respectively, are part of that phenotype. These data and their interpretation form the rationale behind the 72 hour “swarm expansion assay” (72H) (13, 14). The 72H assay is simple, with some minor variations between laboratories: spot mutant cells from a shaking liquid culture onto 0.4% (soft), 1.0%, and 1.5% (hard) agar nutrient plates, allow cells to settle as circular swarms, measure initial swarm diameters, incubate at ~30°C for 72 hours, measure final swarm diameters, and report the difference between final and initial diameters as each mutant strain’s “swarm expansion rate.” The expansion rates of mutant swarms on hard and soft agar in comparison to WT serve as proxies for how much A and S motors, respectively, have been affected by the mutation.

One problem with this interpretation is that it somewhat conflates the broader idea of A and S motility with the narrower idea of the A and S motors. The set of genes involved in *M. xanthus* swarm motility includes all the genes involved in building both A and S motors, along with all the genes involved in their regulation. The A and S motors are genetically distinct, but their regulation is not. Aspects of motility, such as cell speed and reversal frequency, are controlled through a complicated network of overlapping signal transduction systems (15–18). Also, there are many mutations that impact swarm motility only indirectly, and the genes involved may have nothing to do with either motility system.

In laboratory experiments, *M. xanthus* swarms are studied on agar while feeding on casitone. In nature, *M. xanthus* evolved to swarm across many different surfaces and feed on a variety of prey. The phenotypic impact of mutation may not manifest under the deliberately simple and stable laboratory conditions of a 72H assay, or the impact may be subtle, or some of its features may seem unrelated or irrelevant. For example, traveling waves of cell density called ripples sometimes appear within an *M. xanthus* swarm, and multicellular projections called flares sometimes protrude from the edge of an expanding swarm (4, 5, 19–23). These phenotypic features sometimes occur during swarm expansion and, like swarm expansion, they require cell movement coordinated through signal transduction; however, these features have never been included as part of any swarm expansion assay.

We have observed and characterized the *M. xanthus* swarming phenotype beyond the experimental conditions of the 72H assay. By exploring new surfaces, nutrient sources, and data recording methods, we identified novel reproducible and quantifiable phenotypic features that can be applied to genome annotation and genotype-to-phenotype mapping. We then tested whether or not these features can be used to distinguish differences between WT and the phenotypes of 50 single gene insertion-disruption mutant strains that are probably not directly part of either the A or S motility systems.

## Results

We performed swarm expansion assays using seven experimental protocols. We used four surfaces: 0.4% agar (0.4%), 1.0% agar (1.0%), 1.5% agar (1.5%), and a lawn of *Escherichia coli* spotted onto 1.5% agar (Prey). We also used two assay protocols to record and measure data. The first protocol was our version of the 72-hour swarm expansion assay, for which we used a ruler to directly measure changes in swarm diameter every 24 hours for three days. The second protocol is novel and was designed to record swarming dynamics. We used ~40X time-lapse microscopy to record the edge of an expanding swarm at one image per minute, and applied Fiji (38) image processing software to measure swarm expansion every five hours for just over one day. We refer to this as 25H conditions. The final measurement under 25H conditions was made one hour longer than the first measurement under 72H conditions so that the five intervals of 25H conditions would be equal. Both 25H and 72H swarm expansion data are reported as a velocity in millimeters per hour. A more detailed description of surfaces and conditions are provided in Materials and Methods. All four surfaces were used under both conditions with one exception; the combination of Prey under 72H conditions (Prey/72H) was not performed because the WT swarm expanded rapidly enough that it would always expand beyond the boundary of the Prey spot well before the end of the assay. Therefore, 72H conditions were deemed too long for the Prey surface.

WT swarms expand radially from the point of inoculation on all surfaces under both conditions. Flares are observed at the edge of swarms only on 1.0% and 1.5% (Figure 1 panels B, C, F, and G, black arrows), while on 0.4% and Prey, WT swarms had a mostly smooth edge (Figure 1 panels A, D, E, and H, red arrows). Ripples were observed only on Prey (Figure 1 panel D), appearing throughout the swarm within the first few hours and continuing until the end of the Prey/25H assay. A ‘zone of predation’ was also observed on Prey (Figure 1 panel H, green arrow), appearing within the first hour between the advancing edge of the *M. xanthus* swarm and the receding edge of the Prey lawn.

**Figure 1.**
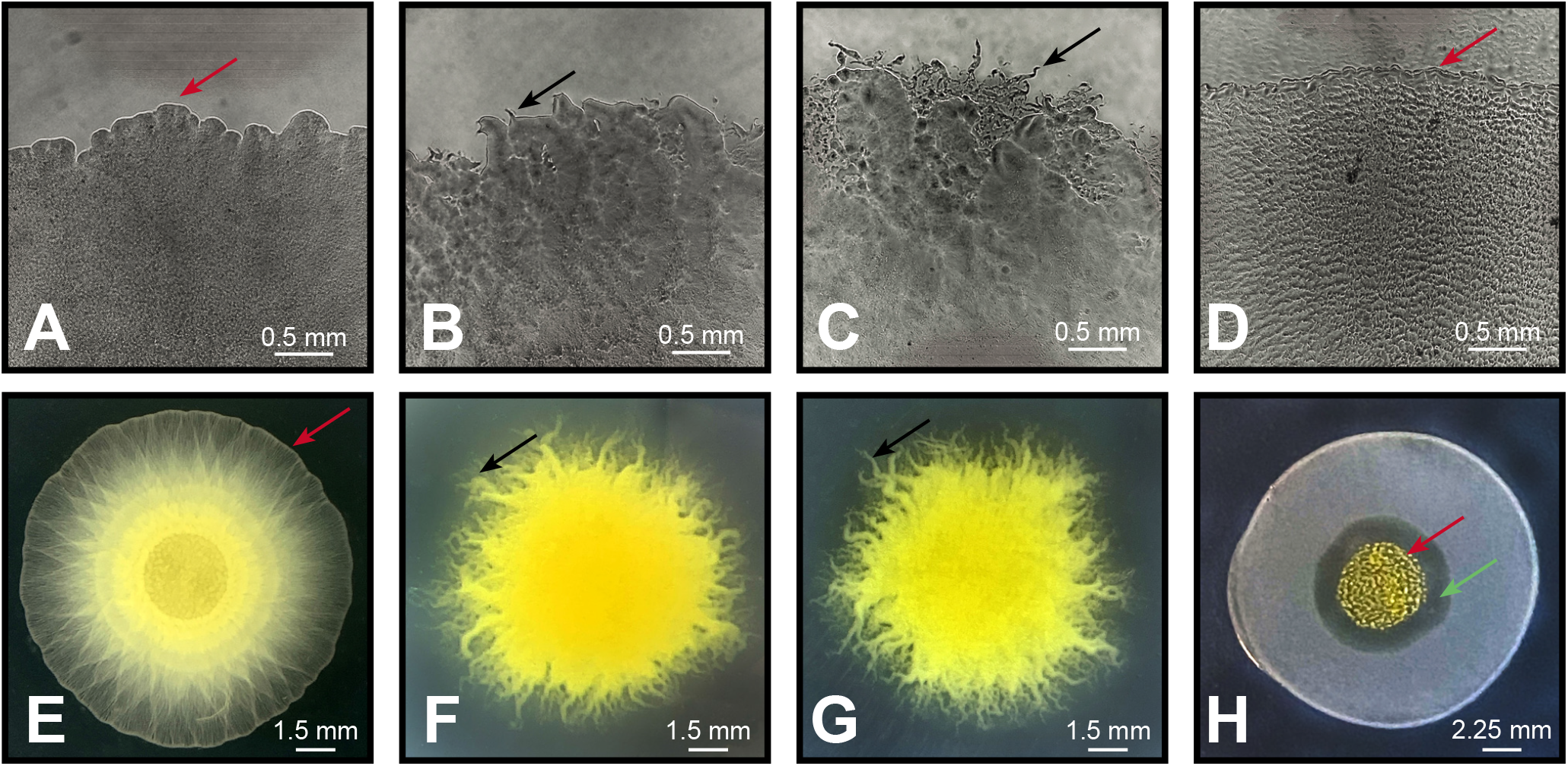
WT *M. xanthus* swarms appear different on different surfaces: 0.4% (A, E), 1.0% (B, F), 1.5% (C, G), Prey (D, H). Top row images (A-D) were the last in a time-lapse image stack taken during a 25H assay (bar = 0.5mm). Image H was also taken after 25 hours of growth (bar = 2.25mm) and images E-G were taken after 72 hours of growth (bar = 1.5mm).

Images from time-lapse microscopy were used to measure swarm expansion at five-hour intervals. Figure 2 shows the progression of a WT swarm on all four surfaces under 25H conditions. To measure swarm expansion on 0.4%, 1.0%, and 1.5%, we measured the position of the swarm edge at four locations within the field of view (Figure 2, first three columns from left). When flares were present on 1.0% and 1.5%, which typically happened at later intervals, measurements were taken at the tip of the furthest flare at each location (for an example, see Figure 2, 1.0% and 1.5%). Using four locations for each timepoint made it possible to account for the irregularity flares caused at the swarm edge; the larger error bars at later time intervals in the graphs represent the impact of flares when measuring expansion. The swarm edge on Prey was the exception. There were almost no edge irregularities at any time point, and error bars were too small to appear in the graphs. Therefore, a single measurement was used to represent swarm expansion on Prey. Also, Prey/25H assays were started with more of the swarm showing within the initial field-of-view, so that the appearance of ripples within the swarm could be observed and recorded.

**Figure 2.**
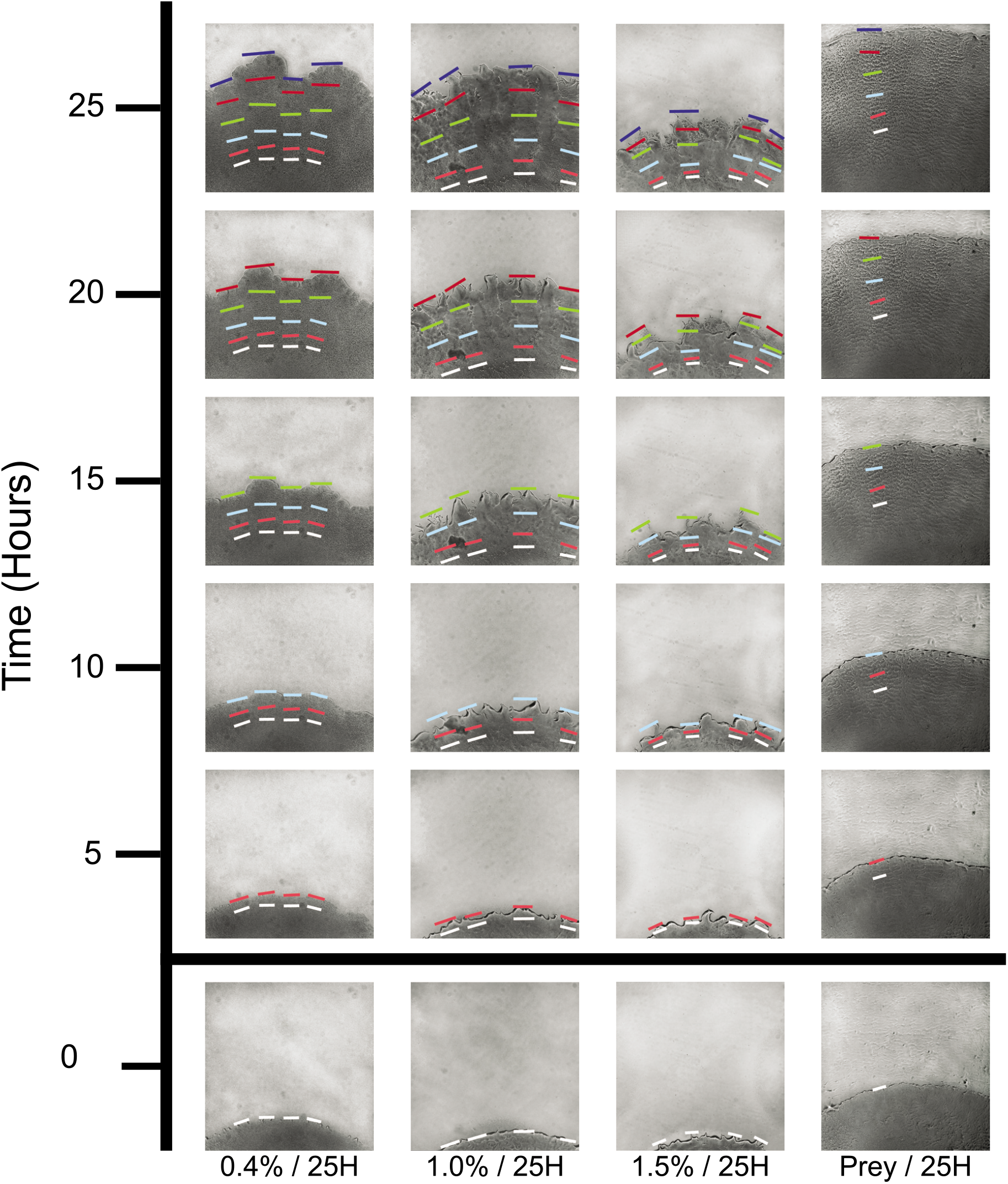
The progression of WT *M. xanthus* swarm expansion on different surfaces. Strains were positioned so only the edge of the swarm was visible on the microscope screen. Swarm expansion was tracked from different positions on the swarm edge. Surfaces used in this study were 0.4% / 25H, 1.0% / 25H, 1.5% / 25H, and Prey / 25H. Colored bars mark the measurements of swarm expansion (white – 0 hours, pink – 5 hours, light blue – 10 hours, green – 15 hours, red – 20 hours, navy – 25 hours).

Graphs representing WT 25H expansion rates on all four surfaces are presented in Figure 3, together with calculated velocities and representative swarm images (Figure 3, first column on left). We also performed the same set of four 25H assays on 50 single gene insertion-disruption mutant strains (Table 1). The disrupted genes included 34 transcriptional regulators from three major families (ECF sigma factors, One Component regulators, NtrC-like activators) and 16 ABC transporters. This assortment was chosen for the following reasons: all four gene families are well represented in the *M. xanthus* genome and many had been the focus of prior research; several had been at least partially characterized (24–31) in *M. xanthus* and were found to have a significant impact on phenotype; and none were deemed likely to be directly involved in either A or S motors based on homology. Figure 3 includes data from some of the mutant strains with notable phenotypes.

**Figure 3.**
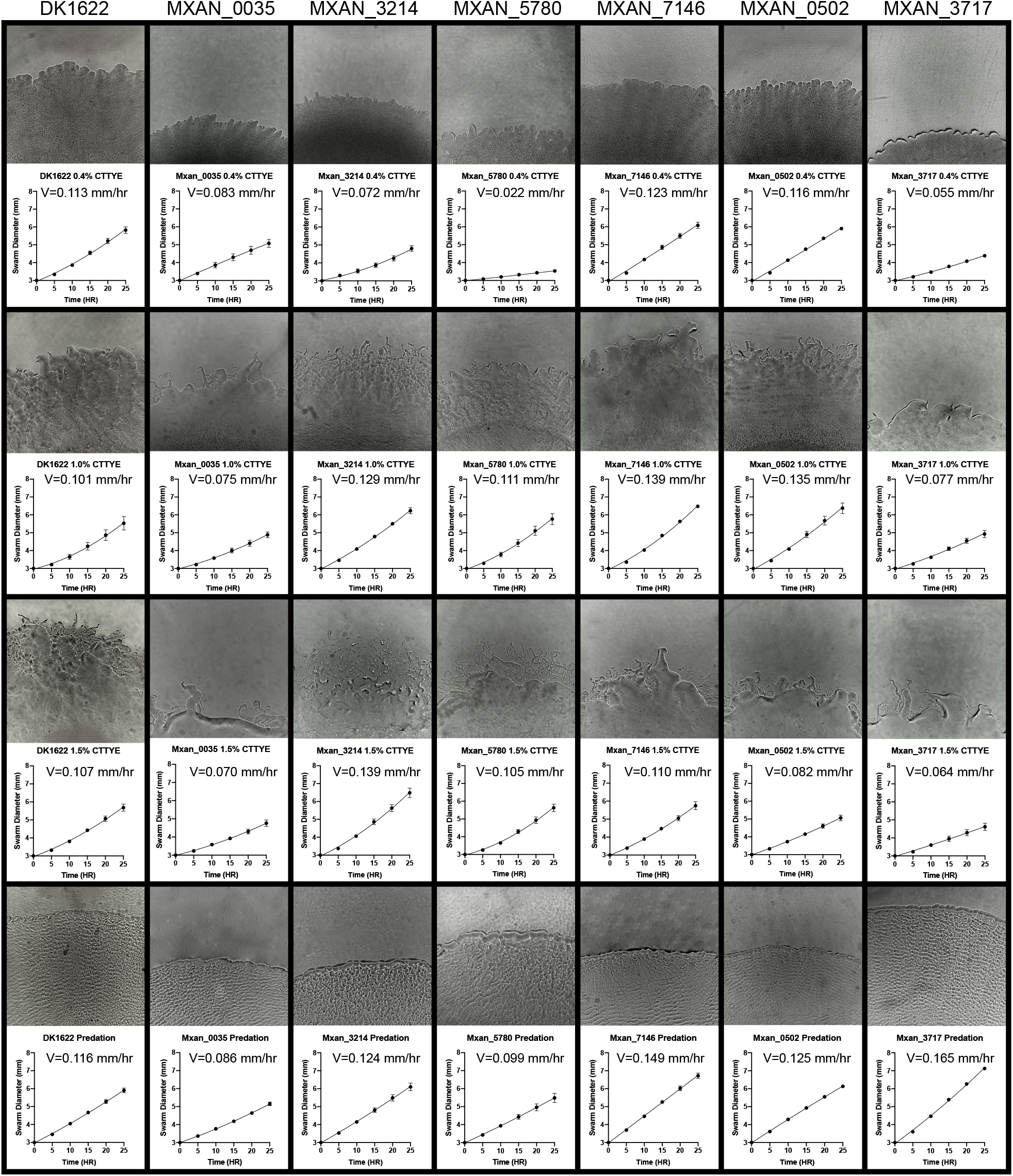
Motility phenotypes were rarely uniform across all assays. While some strains (Mxan_0035) displayed a uniform phenotype, many showed a distinct gain/loss of function on one (or two) surface types (Mxan_3214, Mxan_5780, Mxan_7146, Mxan_0502, and Mxan_3717).

**Table 1.**
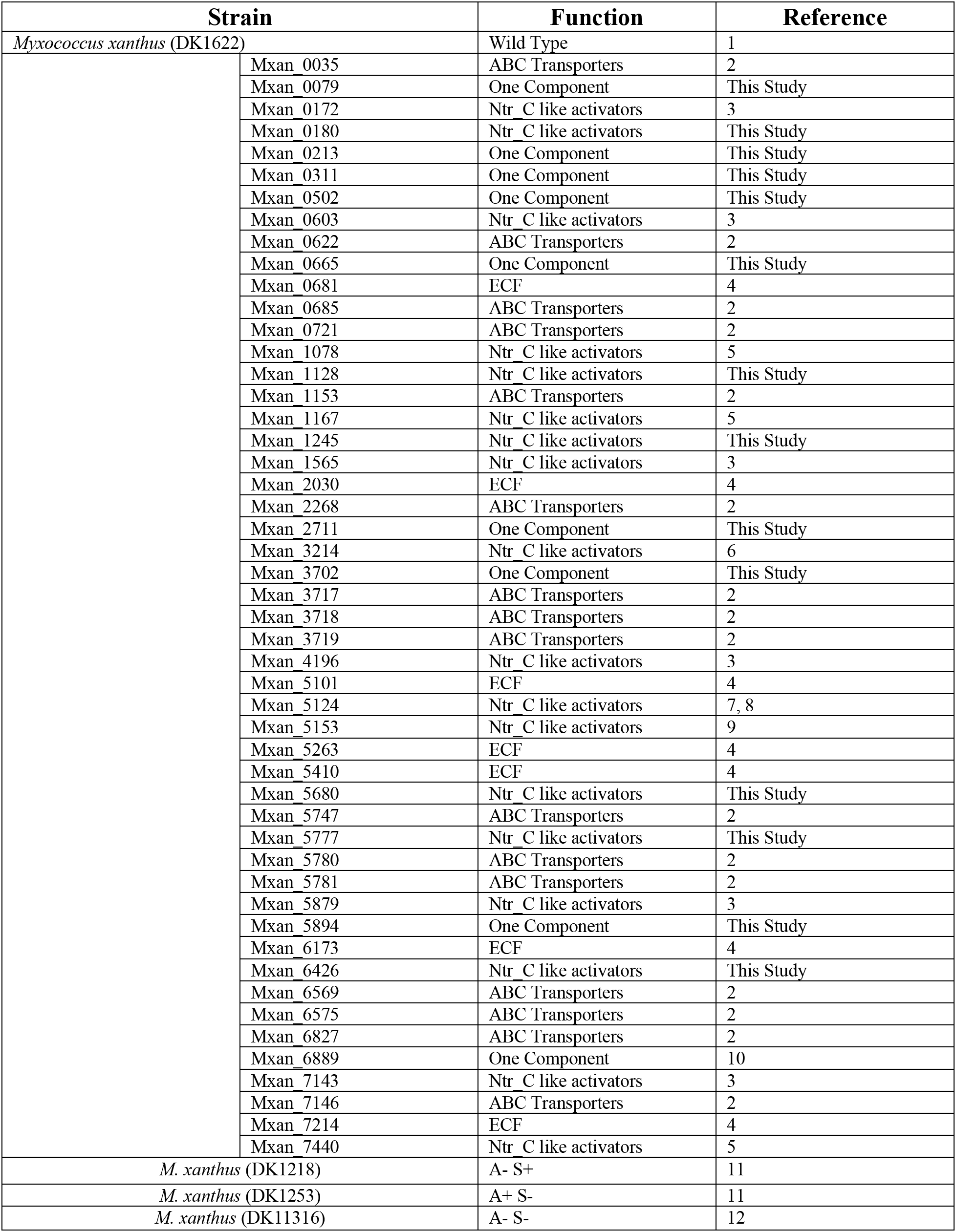

| **Strain** | **Function** | **Reference** |
| --- | --- | --- |
| *Myxococcus xanthus* (DK1622) | Wild Type | 1 |
| Mxan 0035 | ABC Transporters | 2 |
| Mxan 0079 | One Component | This Study |
| Mxan 0172 | Ntr C like activators | 3 |
| Mxan 0180 | Ntr C like activators | This Study |
| Mxan 0213 | One Component | This Study |
| Mxan 0311 | One Component | This Study |
| Mxan 0502 | One Component | This Study |
| Mxan 0603 | Ntr C like activators | 3 |
| Mxan 0622 | ABC Transporters | 2 |
| Mxan 0665 | One Component | This Study |
| Mxan 0681 | ECF | 4 |
| Mxan 0685 | ABC Transporters | 2 |
| Mxan 0721 | ABC Transporters | 2 |
| Mxan 1078 | Ntr C like activators | 5 |
| Mxan 1128 | Ntr C like activators | This Study |
| Mxan 1153 | ABC Transporters | 2 |
| Mxan 1167 | Ntr C like activators | 5 |
| Mxan 1245 | Ntr C like activators | This Study |
| Mxan 1565 | Ntr C like activators | 3 |
| Mxan 2030 | ECF | 4 |
| Mxan 2268 | ABC Transporters | 2 |
| Mxan 2711 | One Component | This Study |
| Mxan 3214 | Ntr C like activators | 6 |
| Mxan 3702 | One Component | This Study |
| Mxan 3717 | ABC Transporters | 2 |
| Mxan 3718 | ABC Transporters | 2 |
| Mxan 3719 | ABC Transporters | 2 |
| Mxan 4196 | Ntr C like activators | 3 |
| Mxan 5101 | ECF | 4 |
| Mxan 5124 | Ntr C like activators | 7, 8 |
| Mxan 5153 | Ntr C like activators | 9 |
| Mxan 5263 | ECF | 4 |
| Mxan 5410 | ECF | 4 |
| Mxan 5680 | Ntr C like activators | This Study |
| Mxan 5747 | ABC Transporters | 2 |
| Mxan 5777 | Ntr C like activators | This Study |
| Mxan 5780 | ABC Transporters | 2 |
| Mxan 5781 | ABC Transporters | 2 |
| Mxan 5879 | Ntr C like activators | 3 |
| Mxan 5894 | One Component | This Study |
| Mxan 6173 | ECF | 4 |
| Mxan 6426 | Ntr C like activators | This Study |
| Mxan 6569 | ABC Transporters | 2 |
| Mxan 6575 | ABC Transporters | 2 |
| Mxan 6827 | ABC Transporters | 2 |
| Mxan 6889 | One Component | 10 |
| Mxan 7143 | Ntr C like activators | 3 |
| Mxan 7146 | ABC Transporters | 2 |
| Mxan 7214 | ECF | 4 |
| Mxan 7440 | Ntr C like activators | 5 |
| *M. xanthus* (DK1218) | A- S+ | 11 |
| *M. xanthus* (DK1253) | A+ S- | 11 |
| *M. xanthus* (DK11316) | A- S- | 12 |

While most mutant strains’ swarm expansion rates were similar to WT, some swarmed slower than WT on all substrates. For example, Mxan_0035 (Figure 3, second column from left) was slower than WT under all conditions. Nevertheless, it produced flares on both 1.0% and 1.5% and ripples on Prey, indicating that the cells were actively moving and forming patterns even though the swarm expanded slowly. While no strains were faster than WT on all agar surfaces, several did swarm faster than WT on at least two. For example, Mxan_3214 (Figure 3, third column from left) swarmed faster than WT on 1.0% and 1.5%. The expansion rate of some strains differed from WT on only one concentration of agar. For example, Mxan_5780 (Figure 3, fourth column from left) was slower than WT only on 0.4%, while Mxan_7146 (Figure 3, fifth column from left) was faster only on 1.0% (and Prey). The rate of Mxan_0502 (Figure 3, sixth column from left) had the greatest variability with respect to agar surfaces, expanding like WT on 0.4% (and Prey), faster than WT on 1.0%, and slower than WT on 1.5%. Finally, several strains displayed a different phenotype on Prey than on any agar concentration. For example, Mxan_3717 (Figure 3, seventh column from left) was slower than WT on all three agar surfaces but was significantly faster than WT on Prey.

We also performed 72H assays on all three agar surfaces for WT and each of the 50 mutant strains, and then compared all swarm expansion rates for all assay conditions. A multiple Spearman correlation graph (36) showing each assay combination reveals positive correlations in every case (Figure 4), although the degree of correlation varies between surfaces and protocols. While discussing the interpretation of correlation data, we will apply the following ‘rule of thumb’ estimates and terms: 0.9-1.0 – ‘very high’, 0.7-0.9 – ‘high’, 0.5-0.7 – ‘moderate’, 0.3-0.5 – ‘low’, 0.0-0.3 – ‘negligible’. Most notably, there were high correlations between the three agar surfaces under 72H conditions (0.86, 0.89, 0.89). Surface correlations under 25H conditions were high for 1.0% compared to 1.5% (0.82), but moderate for 0.4% compared to 1.0% (0.48) and 0.4% compared to 1.5% (0.46). Correlation between 25H and 72H conditions for the same agar surface were moderate for 0.4% and 1.5% (0.56, 0.66) and high for 1.0% (0.73).

**Figure 4.**
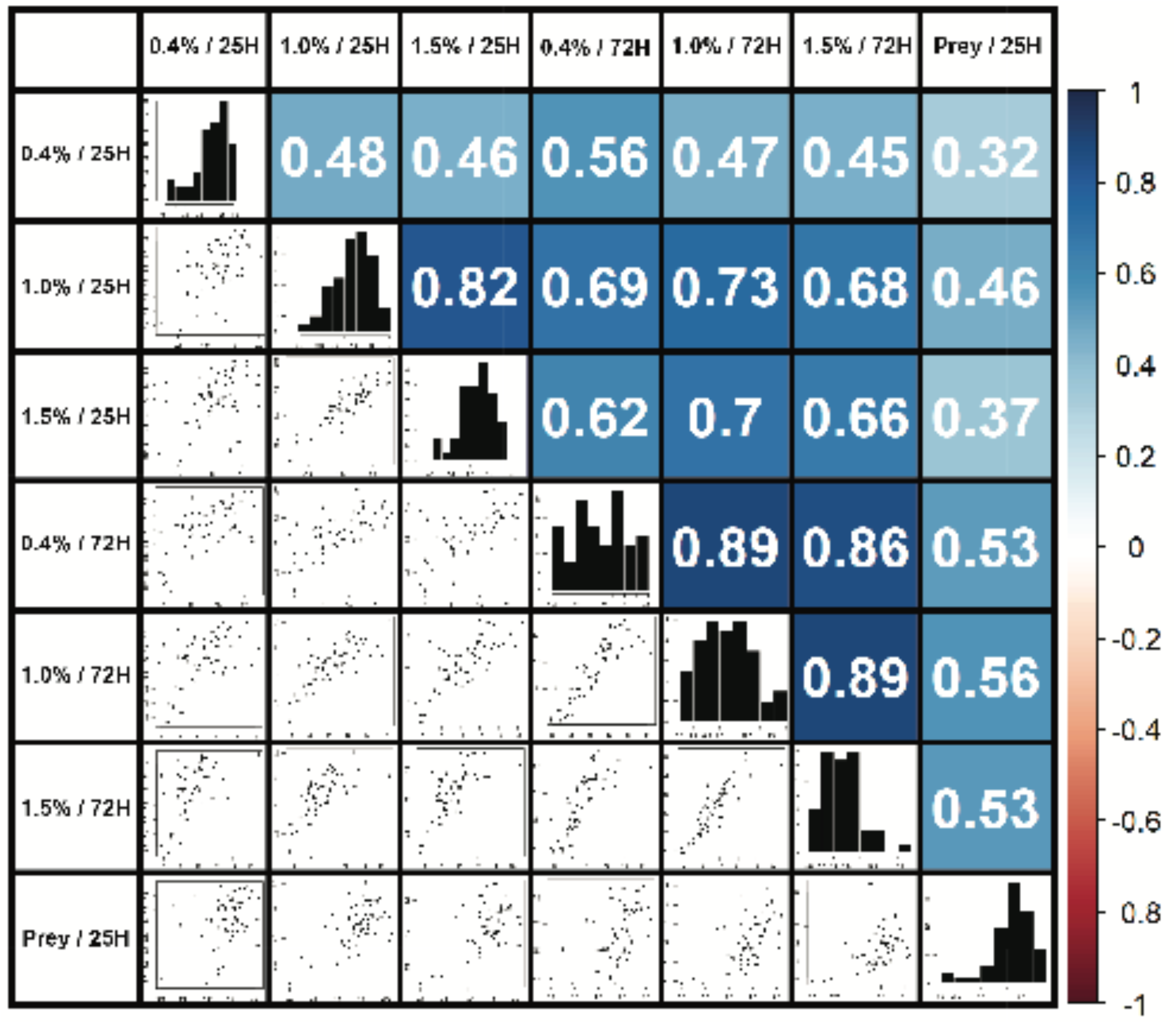
Motility assays are correlated, but not equally. Scatter plots (left side) show pairwise comparisons of expansion data for all assay combinations of swarm motility data for WT and mutant strains. Bar graphs (center diagonal) are histograms showing the distribution of swarm expansion data for each assay. Numbers (right side) show the Spearman’s rank correlation coefficient for each pairwise assay comparison.

Swarm velocity on Prey did not show high correlation with any agar surface using either condition. Correlation was low between Prey and all agar surfaces (0.32, 0.46, 0.37) using 25H conditions, and moderate between Prey and all agar surfaces using 72H conditions (0.53, 0.56, 0.53).

Three older strains (DK1218, DK1253, and DK11316) were included as reference controls to confirm that the results of this study match those of prior work, including the initial reports relating the different impact of agar concentration on A and S motility (4,5). DK1218 (*cglB2*) is defective in A motility (A-S+), DK1253 (*tgl-1*) is defective in S motility (A+S-) (4,5), and DK11316 is nonmotile (AS-) (pilA::Tc^r^ΔcglB) (32). The relative swarm expansion rates for these controls match prior published results, (Figure 5, colored triangles). The three control strains expand more slowly than WT on all surfaces using both 25H and 72H conditions; DK1218 is faster than DK1253 on 0.4%, and DK1253 is faster than DK1218 on 1.5%. Only DK1218 formed flares, and only on 1.0% and 1.5%, and none of these reference control strains formed ripples on Prey, suggesting that both A an S motility are required for rippling.

**Figure 5.**
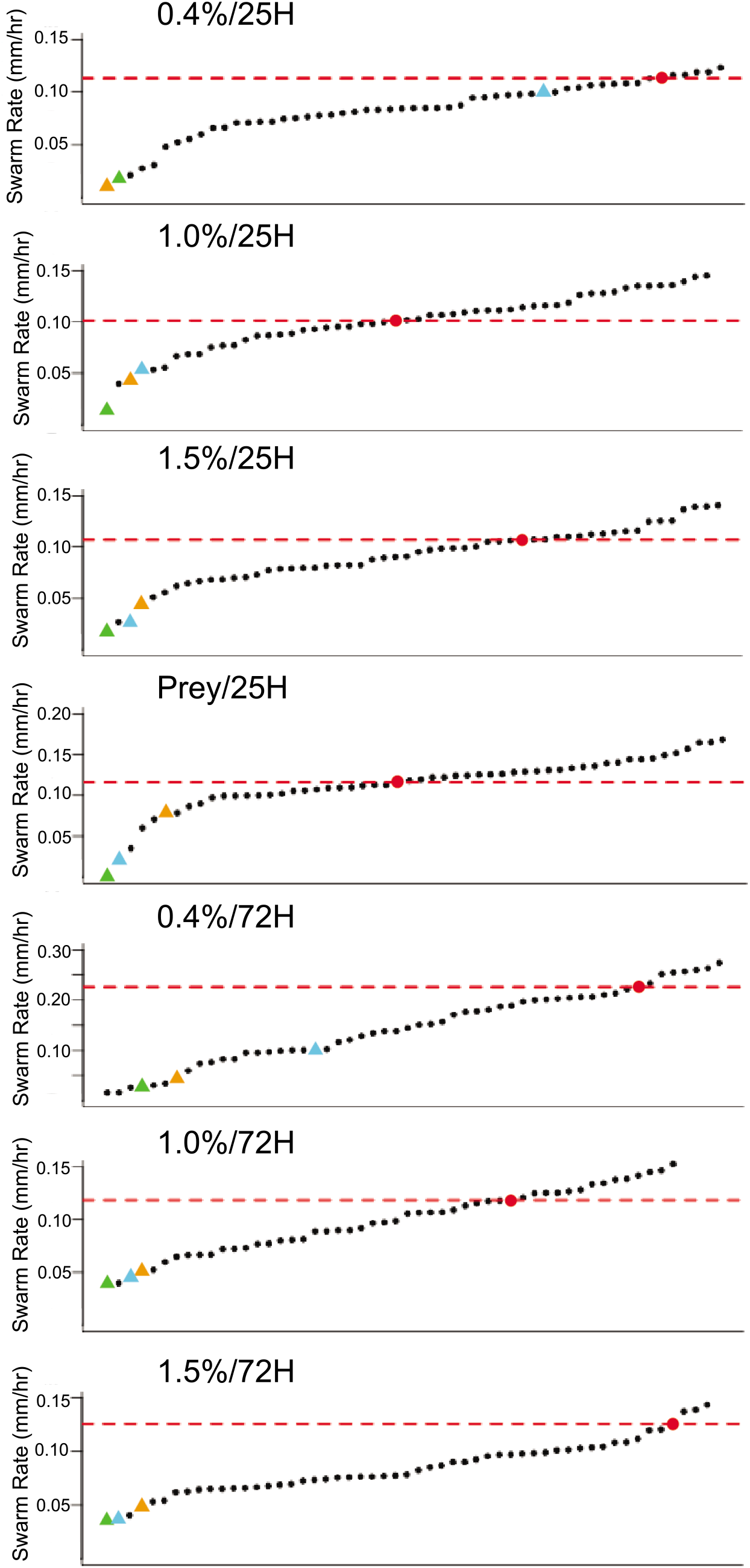
WT and mutant strain *M. xanthus* swarm expansion rates form a continuous distribution. Each graph shows the swarm expansion rates arranged from slowest (left) to fastest (right). The expansion rate of WT (red dot) is marked with a dotted red line. Reference mutant strains are marked as colored triangles: green - DK11316, blue - DK1218, and orange - DK1253. The rates of all 50 mutant strains are in Table S2.

Expansion rates for WT, the reference controls, and the 50 mutant strains were combined and displayed in increasing order according to their expansion rates for all seven assays (Figure 5). The graphs have at least three notable features: (1) all display a continuous distribution; (2) the majority of strains fall within a confidence interval one standard deviation about the mean; and (3) WT is within this confidence interval for 4 of the 7 graphs and, and for the remaining cases (0.4%/25H, 0.4%/72H, 1.5%/72H) WT is above it.

Time-lapse image data from WT and the 50 mutant strains also have several notable features regarding ripples and flares. Rippling on Prey proved to be a robust phenotype, observed in 49 of the mutant strains. The only strain that appeared not to ripple, Mxan_7440, showed little swarm expansion on any surface using either protocol, although several strains that had similarly low expansion rates did ripple on Prey (Mxan_6827, Mxan_7143, Mxan_0035, Mxan_1078, etc.). The ripples of several mutant strains appeared noticeably different from WT with respect to their intensity, speed, wavelength, and other discernable features, but these differences proved difficult to quantify (Figure 6, top row). For example, a swarm of Mxan_0213, despite expanding faster than WT on Prey, appeared to produce smaller slower ripples. A swarm of Mxan_1128 also appeared to have slower ripples throughout predation, despite having an expansion rate similar to WT.

**Figure 6.**
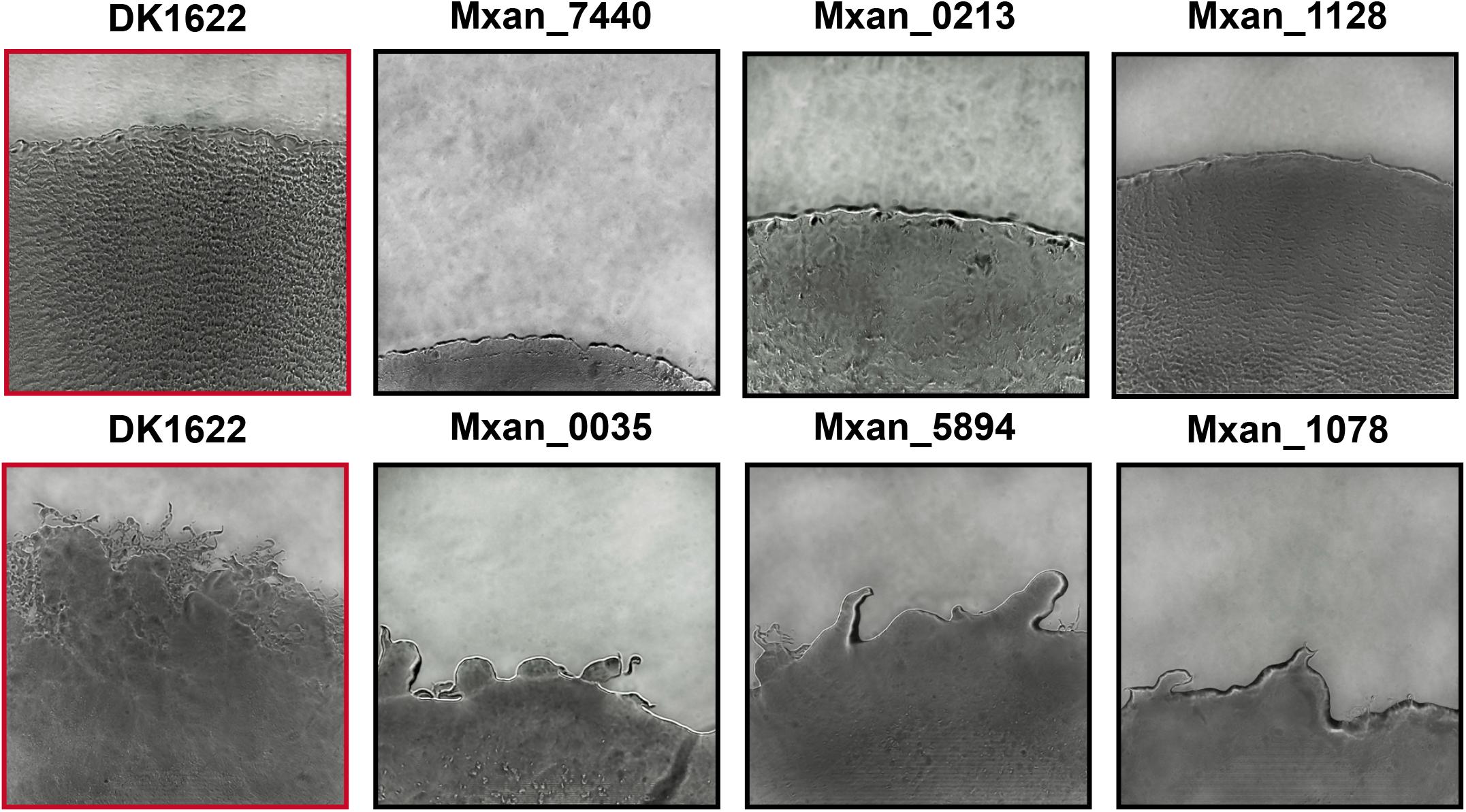
Some mutant swarm phenotypes are qualitatively different from WT. Images are swarms on Prey (top row) or 1.5% (bottom row). WT is shown on Prey (top left red box) and 1.5% (bottom left red box). Mxan_7440 was the only strain to display no ripples when plated on *E. coli*. Mxan_0213 and Mxan_1128, despite expanding at least as quickly as WT, formed ripples that appeared to have a longer wavelength (Mxan_0213) or slower velocity (Mxan_1128). Mxan_0035, Mxan_5894, and Mxan_1078 are mutant strains that produce a smaller number of flares that are noticeably different from WT when plated on 1.0% or 1.5% agar.

Flares were observed in most mutant strains, although they proved to be less robust than ripples. Like ripples, the flares of several mutant strains appear different from WT with respect to discernable features, such as length, width, number, curvature, and branching, but these also proved difficult to quantify (Figure 6, bottom row). However, some strains produced an irregular edge, but no distinct protrusions that we would call flares on harder (1.0% and 1.5%) agar surfaces, and these we were able to distinguish from WT. Mxan_0035, Mxan_5894, and Mxan_1078 produced these edges on 1.0% and 1.5% agar. Their expansion rates were mostly slower than WT, except for Mxan_5894 on 1.0% agar, which was similar to WT.

## Discussion

An *M. xanthus* swarm’s expansion rate is a single measurement that does not fully represent the complicated swarming phenotype. Swarms expand at different rates on 0.4% and 1.5% agar, and they look different in other ways that are hard to describe and harder to quantify. These observations are true for both WT and mutant strains. By varying the laboratory environment beyond standard assay conditions and by collecting some additional data across a set of mutant strains, we were able to identify and characterize several new features of the *M. xanthus* swarming phenotype.

The 72H assays on 0.4%, 1.0%, and 1.5% are closest to the standard swarm expansion assay conditions described dozens of times. The relative expansion rates of the 50 mutant strains provides additional information about the WT phenotype. WT was close to the fastest on 0.4% agar, but on 1.0% and 1.5% it was near the mean. This observation is important because it may indicate the probability of any future mutant strains being faster or slower than WT on different agar concentrations. On 0.4% it is much more likely that mutant strains will be slower than WT, but on 1.0% or 1.5% they could be slower or faster with roughly equal probability. If we hypothesize that the continuous distributions of expansion rates for WT and these 50 mutant strains under 72H conditions represent a very preliminary approximation of the distributions for all mutant strains under 72H conditions, then it would be reasonable to conclude that expansion rate is not a trait that has been optimized in WT, especially for higher agar concentrations.

There was a high degree of correlation between 0.4%, 1.0%, and 1.5% 72H assays across all mutant strains. At nearly 0.9, it indicates that measuring the 72H expansion rate at different agar concentrations may not provide meaningful discriminatory power above measuring it at just one agar concentration. Surprisingly, 0.4%, 1.0%, and 1.5% 25H assays displayed only modest correlation when compared to their 72H counterparts. It seems implausible that swarm expansion rates would be closely correlated under 72H conditions for all three agar concentrations, but only moderately correlated between 72H and 25H assays at the same agar concentration. After all, both 25H and 72H assays are presumably measuring the same phenotype in two different ways. The high degree of correlation between 0.4%, 1.0%, and 1.5% also appears limited to the 72H assays, as the three agar concentrations display only modest correlation under 25H conditions.

Why does correlation between agar concentrations increase between 25 and 72 hours? One explanation is that, for many of the mutant strains, swarm expansion does not occur at a constant rate. A comparison of the distributions for 0.4%/25H and 0.4%/72H plots in Figure 5 reveal that enough strains accelerate on 0.4% agar between 25 and 72 hours that it forces a change in scale on the Y-axis between the 0.4%/25H and 0.4%/72H plots. Examples of accelerating strains are Mxan_0172, Mxan_0213, Mxan_0665, Mxan_5101, Mxan_5780, and Mxan_6575. Although fewer in number, it is important to note that a few strains actually decelerate on 0.4% agar between 25 and 72 hours, such as Mxan_0685 and Mxan_1245. Acceleration and deceleration are also not limited to 0.4% agar. One notable strain on 1.5% agar, Mxan_0172, accelerates from slower to much faster than WT between 25 and 72 hours. On the 1.5%/72H plot, Mxan_0172 is by far the fastest strain, appearing as an outlier on the right side of the distribution.

Why would swarm expansion rates change between 25 and 72 hours so that 72H data would display a higher degree of correlation than 25H data for all three agar concentrations? At this point, hypotheses that attempt to explain the biological underpinnings of this phenomenon must be based on scant evidence. Perhaps some accelerating mutant strains synthesize exopolysaccharide (EPS) more slowly than WT. Perhaps some decelerating mutant strains have growth and division rates that are slower than WT. Perhaps the manifestation of mutant phenotypes requires different amounts of time on different surfaces, and 72 hours is sufficient to affect swarms on all surfaces to a roughly equal degree. Perhaps this is why only the 72H data displays a high degree of correlation. All or none of this might be correct but determining the biological cause of the mutant phenotypes is not the purpose of this study. We are trying to assess the discriminatory power of different features of the swarm motility phenotype to make meaningful genotype-to-phenotype associations. The purpose of this study is to develop assays that provide greater discriminatory power because they measure a larger set of distinct independent variables that are under genetic control.

From this perspective, prey as a surface is more useful than hard or soft agar at discriminating between mutant swarm phenotypes because its correlation to any of the agar surfaces was only modest. *M. xanthus* is a saprophytic predator; a swarm secretes enzymes and antibiotics to lyse prey extracellularly, then consumes the released cellular contents as it expands (33). This feeding behavior can be observed under a microscope, where a small cell-free zone develops between the expanding outer edge of a swarm and the receding edge of the prey population (34). This zone of predation was observed on WT and all mutant strains used in this study. It would therefore be reasonable to postulate that the surface on which the swarm is expanding is actually the 1.5% agar below the prey bacteria, and that prey bacteria represent a nutrient source and not a surface. This may be true, but *M. xanthus* swarms on prey do not look like swarms on 1.5% agar. The expansion rate of a WT swarm on prey is faster than on 1.5%, and both WT and all mutant strains lack flares on prey. In these ways swarms on prey appear closer to swarms on 0.4% agar than on 1.5% agar, and the ubiquitous rippling on prey makes it different from both.

Ripples and flares are both phenotypic features of swarm motility, but their purpose remains unknown. Flares clearly aren’t required to move quickly over surfaces, since swarm expansion on Prey and 0.4% agar was mostly faster than on 1.0% or 1.5%, and no strains produced flares on either surface, including WT. Ripples also don’t seem to be related to swarm expansion on Prey, since strains that expanded slowly and strains that expanded quickly all produced ripples (20, 23). Some mutant strains produce ripples and flares that are discernably different from WT, which indicates that these phenotypic features are under genetic control. This alone makes them useful in making genotype-to-phenotype associations.

## Materials and Methods

### Bacterial strains and growth conditions

WT *M. xanthus* (strain DK1622), 50 single gene insertion-disruption mutants, DK1218, DK1253, DK11316, and *E. coli* (K12) were used in this study (Table 1). WT *M. xanthus* was grown in CTTYE media (1.0% Casitone, 0.5% yeast extract, 10mM Tris-HCl (pH 8.0), 1mM KH_2_PO_4_, and 8mM MgSO_4_) or on petri dishes containing CTTYE broth and either 0.4, 1.0 or 1.5% agar. Both plates and liquid cultures were incubated at 32°C. Mutant insertion-disruption *M. xanthus* strains were grown in CTTYE supplemented with kanamycin (40μg/mL) as a selective agent. *E. coli* was grown in LB broth (Sigma, Cat#: L3022) and LB broth with 1.5% agar. Predation assays were performed using TPM plates (10mM Tris-HCl (pH 7.6), 1mM KH_2_PO_4_, 8mM MgSO_4_) with 1.5% agar.

All motility assays were performed using bacteria under these growth conditions. Assays are named using the convention: *Surface substrate (0.4%, 1.0%, or 1.5% agar, or Prey) / Time and assay conditions (25H or 72H)*

### 72-hour Swarm Expansion Assay

Both WT *M. xanthus* and mutant strains were inoculated in CTTYE broth (40μg/mL Kanamycin added to mutant cultures) and incubated overnight at 32°C with agitation (~300 rpm). Once cell density was ~100 Klett (~5×10^9^ cells/mL), cells were harvested and resuspended to 1,000 Klett in CTTYE. Five 2μL spots were then placed on CTTYE plates of varying agar concentrations (0.4%, 1.0%, and 1.5%). The plates were incubated at 32°C for 72 hours and measurements (in 1mm increments) of each spot were taken every 24 hours using a ruler.

### 25-Hour Motility Assay

Both WT *M. xanthus* and mutant strains were inoculated in CTTYE broth (40μg/mL Kanamycin added to mutant cultures) and incubated overnight at 32°C with agitation (~300 rpm). Once the cell density was ~100 Klett (~5×10^9^ cells/mL), cells were harvested and resuspended to 1,000 Klett in CTTYE. Five 2μL spots were placed on CTTYE plates of varying agar concentrations (0.4%, 1.0%, and 1.5%). The plates were incubated at 32°C for two hours before being inverted and placed on a 3D-printed brightfield microscope (https://www.thingiverse.com/thing:2672065). The microscope was focused on the edge of one of the 2μL spots so the edge of the spot took up less than ¼ of the screen, leaving the rest of the field of view open in the direction that the spot would move. One image was taken every 60 seconds for 25 hours. Using Fiji (38), the images were compiled into a timelapse movie showing how the strain swarmed over 25 hours.

### Prey Assay

Both WT *M. xanthus* and mutant strains were inoculated in CTTYE broth (40μg/mL Kanamycin added to mutant cultures) and incubated overnight at 32°C with agitation (~300 rpm). Once cell density was ~100 Klett (~5×10^9^ cells/mL), cells were harvested, washed 2X in TPM broth, and resuspended to 1,000 Klett in TPM. *E. coli* cells were grown in LB broth at 32°C with agitation (~300 rpm) and harvested at an OD_600_ of ~0.8. The *E. coli* cells were washed 2X in sterile autoclaved water, resuspended in autoclaved water, and 40μL were spotted in the center of a TPM plate (1.5% agar). Once the spot dried, 2μL of washed and resuspended *M. xanthus* cells were spotted directly on top of the dried *E. coli* spot. The plates were incubated at 32°C for two hours before being inverted and placed on a 3D-printed brightfield microscope (https://www.thingiverse.com/thing:2672065). The microscope was focused on the edge of one of the 2μL *M. xanthus* spots so the edge of the spot took up approximately ¼ of the screen, leaving the rest of the field of view open in the direction that the spot would move. One image was taken every 60 seconds for 25 hours. Using Fiji (38), the images were compiled into a timelapse movie showing how the strain had swarmed over *E. coli* during the 25 hours.

### Data Analysis and Availability

Using images from time-lapse movies, we marked the edge of the spot at times 0-, 5-, 10-, 15-, 20-, and 25-hours using Fiji (38). We measured the distance in pixels between the timepoints and converted the pixels to millimeters. Because the swarm expanded radially, and the initial swarm diameter created by the 2 μL spot was consistently 3mm, we used each measurement to calculate overall swarm diameter at each timepoint: *Swarm Diameter = (Timepoint measurement (mm) x 2) + 3mm*. For each video taken on CTTYE plates, four sets of measurements were taken at different locations along the spot edge in order to account for uneven swarm expansion (flares) on harder surfaces. WT and mutant strains each had at least four replicates for each surface tested (0.4%, 1%, 1.5% and prey). Correlation analysis, comparison, and visualization were done using R Studio (35), corrplot (36), ggplot (37), and Prism 9 version 4.00 for Windows, GraphPad Software, San Diego California USA, www.graphpad.com. All data collected in this study is available in the Supplementary material (Tables S1 and S2).

## Conclusion

For a laboratory model organism with a reasonably long history of molecular genetic studies, such as *M. xanthus,* a set of ‘standard’ phenotype assays emerges over time as a means of relating current experiments to prior work. The utility of these assays therefore depends on their reproducibility, thus consistent protocols and analyses become established over time. Often a narrative will evolve along with an assay, providing a biological explanation for phenotypic data. It can be useful to depart from these assays and narratives through naive genotype-to-phenotype association studies to provide a broader context and fresh perspective on the relationship between an organism’s phenotype and its species’ phenome, as we have done in this study of swarm motility. By tweaking the protocol and analysis of the standard 72 hour swarm expansion assay, we were able to observe and quantify more granular detail about the phenomenon of expansion, e.g., that expanding swarms can change their velocity, that WT *M. xanthus* seems optimized to expand on a softer agar surface, that a harder agar surface seems to favor the formation of flares, that rippling is a robust phenotype on Prey, and that Prey as a nutrient source changes the swarming phenotype in many ways beyond rippling. Although most of these findings appear tangential to the standard A and S motility narratives, they help us to better understand which phenotypes fall within “normal” parameters, and which are outliers that may exist at the edge of the *M. xanthus* phenome.

## Acknowledgements

This work was supported by National Science Foundation grant DBI-1244295 to R.D. Welch. We thank Caitlin McDonough-Goldstein for helpful discussions on statistical analysis and coding as well as Jessica Comstock and Laura Welch for help in editing this manuscript.

